# Comparison of a long-read amplicon sequencing approach to short-read amplicons for microbiome analysis

**DOI:** 10.1101/2025.09.04.674341

**Authors:** Brandon O’Sullivan, Katherine W. Herbst, Alexander H. Hogan, Michele Maltz-Matyschsyk, Justin D. Radolf, David Lawrence, Michael A. Lynes, Juan C. Salazar, Joerg Graf, The Connecticut Children’s COVID Collaborative

## Abstract

Most microbiome studies to date rely on sequencing short amplicons of the 16S rRNA gene on Illumina’s platforms. Because of the short read length, sequences often can be identified reliably only to the family or genus levels. Long read sequencing with whole-length 16S rRNA sequencing can improve taxonomic resolution, but often only to the species level. StrainID is an alternative approach that amplifies a large segment of the ribosomal operon, including the entire 16S rRNA gene, internal transcribed spacer, and a portion of the 23S rRNA gene. This longer amplicon is designed to allow ribotype-level classification. Although studies have demonstrated the utility of StrainID for several sample types, it has not yet been validated for saliva. Here, we compared the performance of StrainID to short read amplicons with saliva samples as well as a synthetic mock DNA community and human and mouse fecal samples. Short reads were amplified with primer pairs appropriate for the corresponding sample type, and were classified with two different taxonomic databases. For both saliva and fecal samples, we found that StrainID performed similarly to short reads overall and demonstrated a key benefit with phylogenetic-based beta diversity tests and taxonomic classification. Our results further build on establishing StrainID as a valid method and specifically validate its use with saliva samples.

**Importance:** The interpretation of microbiome composition studies is highly dependent on the methodologies chosen during experimental design, which affects factors such as resolution, throughput, cost, and accuracy. StrainID is an approach that can improve resolution while maintaining high-throughput and similar costs to short-read sequencing. The salivary microbiome represents a diverse community of microbes with links to a variety of health conditions and disease states. Closely related strains of bacteria can have drastically different effects on their host. Establishing StrainID as a valid approach for studying the salivary microbiome opens avenues for research that improve upon alternative methods by increasing sensitivity and accuracy compared to traditional short read approaches.

## Introduction

Next-generation sequencing has revolutionized microbiome research, enabling rapid, highly accurate characterization of hundreds of microbial communities simultaneously (1, 2). However, most existing research has relied on short-read amplicon sequencing approaches, focusing on small hypervariable regions within the 16S ribosomal RNA (rRNA) gene and, consequently, limiting taxonomic resolution to genus-level or higher classifications (3). Careful selection of primers for a specific region of the 16S rRNA gene or other genes can improve resolution for certain sample types by focusing on regions that are most variable for the abundant taxa in that environment, although this does not overcome all limitations of this approach (4). Shotgun, or non-targeted metagenomics, offers a potential solution to this problem by sequencing random fragments of all DNA present in a sample. This approach provides much more data on the community, but it is significantly more expensive and computationally taxing than short-read amplicon sequencing and generally requires both higher quality and quantities of DNA (5, 6). One potential outcome of using this approach is greater insights into the metabolic potential of the abundant members of the microbiome at the cost of not obtaining information about less abundant microorganisms (7). These low-abundance organisms can play important roles in determining the composition and function of microbiomes (8). More recently, long-read sequencing technologies, such as those done with PacBio or Oxford Nanopore instruments, have the potential to strike a middle ground by amplifying and sequencing the entire 16S rRNA gene. This typically increases resolution to species-level classification, representing a major improvement over short-read amplicon (SRA) sequencing, but fails to identify strains as metagenomics can (3). Strains from the same species can differ in metabolic function (8), toxin production (9), and antibiotic resistance (10), each with their own potential effects on the host. As such, choosing a technology that allows one to analyze a large enough cohort to detect compositional differences between microbiomes and with sufficient resolution at the sequence level to reveal potential differences is key for the success of clinical studies.

StrainID, an approach commercialized by Intus Biosciences, attempts to improve on whole-length 16S rRNA sequencing at a similar cost by including more of the ribosomal RNA operon. Compared to the approximately 1,500 bp length of the entire 16S rRNA gene, StrainID uses custom primers to produce an amplicon nearly 2,500 bp in length that includes the internal transcribed spacer region (ITS) and a portion of the 23S rRNA gene (Figure 1A) (11, 12). Combined with the custom Athena database, consisting of high quality rRNA operon sequences from published genomes, StrainID is reported to provide strain-level classifications (11). Because strains can be defined by differences in other key genes without any changes in the rRNA operon sequence, these classifications are more accurately called ribotypes instead (13). Herein, we refer to these ribotypes as strains for consistency with their identification by classifiers. Nonetheless, ribotype-level classification represents a substantial improvement over species-level classification.

**Figure 1:**
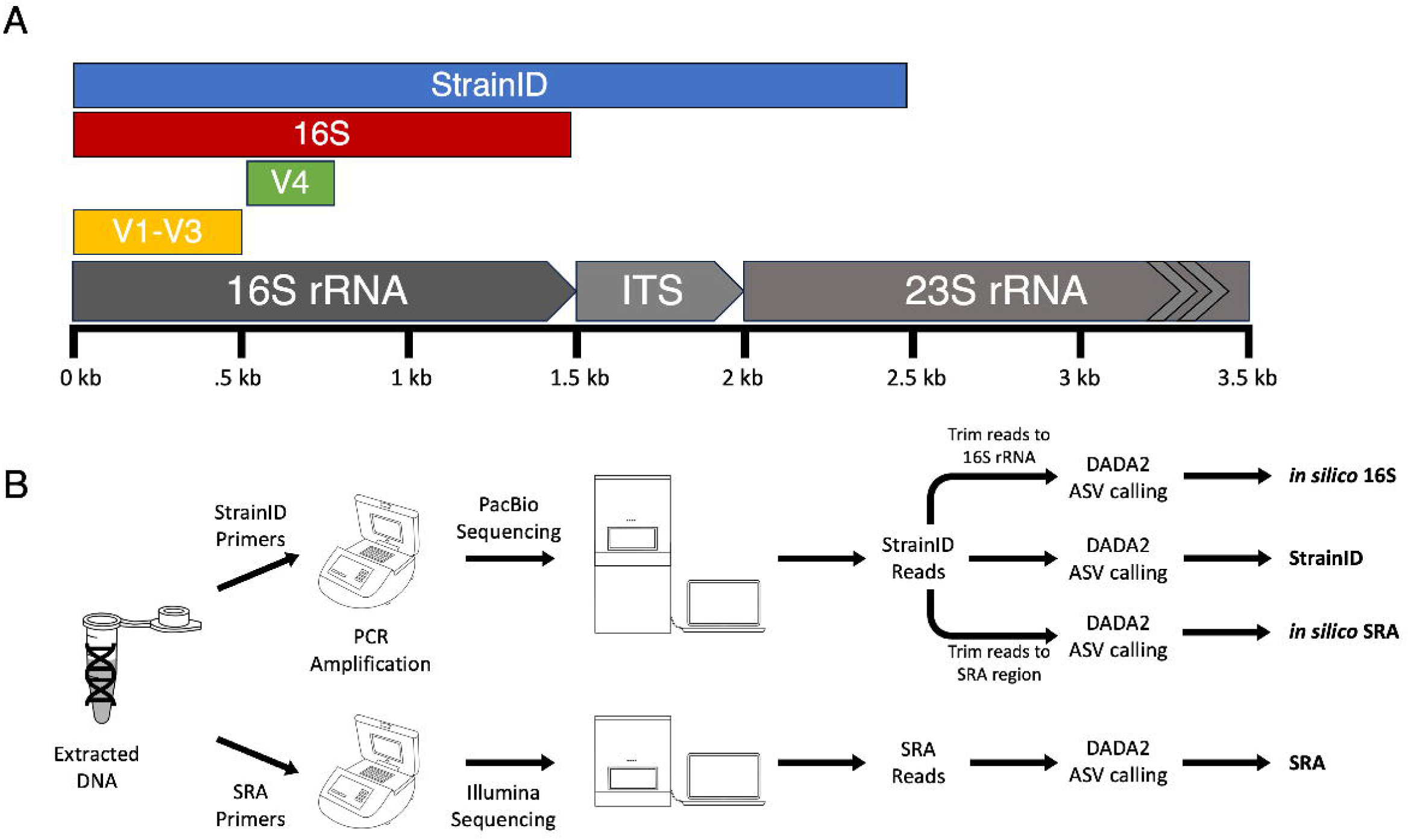
Experimental design. (A) Size and position of the amplicons used in this study relative to a standard ribosomal RNA operon. (B) Sample processing pipeline for downstream data analysis. Every sample was processed with all methods to generate four total datasets.

Most bacteria carry multiple copies of the ribosomal operon in their genome, and these copies frequently have different sequences, especially in the ITS region, often encoding transfer RNA (tRNA) (14). Past studies have shown that the StrainID approach can be used to identify distinct patterns of these sequences that co-occur, generating a “fingerprint” that can be used to identify and track specific bacteria that share the same set of rRNA sequences, or ribotypes, across samples (11, 12). To date, StrainID has been validated for several sample types, including fecal, human milk, and aquaculture water and swabs (11, 12, 15, 16). However, this approach has not yet been validated for the analysis of human saliva.

The human oral microbiome, the second largest and most diverse bacterial community in the human body, contains numerous medically-significant bacteria (17, 18). The oral microbiome is temporally stable and can be influenced by environmental factors (19). Additionally, due to the prevalence of biofilms, the oral microbiome tends to be less susceptible than the gut microbiome to disruption by antibiotics (20). Changes in the relative abundances of certain bacterial taxa in saliva have been correlated with various systemic pathologies including cardiovascular disease, rheumatoid arthritis, and Alzheimer’s disease (18). These findings, combined with the less invasive nature of saliva collection, raises the possibility that saliva could replace blood for diagnostic tests in some contexts (21, 22). However, approaches to identify these signals must be sensitive and specific. With short read approaches, the V1-V3 hypervariable regions represent the section of the 16S rRNA gene that provides the greatest ability to resolve taxa associated with saliva, especially *Streptococci* (23, 24). For Illumina sequencing, the length of this amplicon requires the use of Illumina’s 600-cycle kit which generates paired-end 300 bp reads. The chemistry used in Illumina sequencing is associated with a decrease in read quality as the length increases, especially in the reverse read (25). This problem is far more pronounced in the 600-cycle kit compared to even the 500-cycle kit (26). The decrease in quality can result in paired-end reads failing to join due to insufficient overlap, and reads with low quality may have to be discarded for failing quality filtering, especially when using amplicon sequence variants (ASVs). In addition, longer reads that contain more variable sites than short reads can potentially provide a greater sensitivity in distinguishing different cohorts especially when using phylogenetic distance-based metrics (27). Alternative sequencing technologies might overcome these shortcomings.

Herein, we sought to evaluate the performance of StrainID with human saliva samples to test both its accuracy and robustness in comparison to V1-V3 amplicon sequencing. To increase confidence in our findings and their applicability, we confirmed the performance of StrainID with a mock DNA community representing diverse bacteria, including Gram-positive and Gram-negative organisms, and with fecal samples from humans and mice. We selected mock communities and fecal samples for specific reasons. Mock communities have been established as a way to verify the success of sequencing runs and estimate biases (28). The fecal microbiome is the largest and most diverse bacterial community in the human body, and also the most extensively studied.

The V1-V3 regions are often used to study the salivary microbiome, whereas amplicons including the V4 region are commonly used for fecal microbiome studies (23, 29). As such, we used saliva samples to compare StrainID specifically against V1-V3 amplicons, and fecal samples to compare StrainID to V4 amplicons (Figure 1B). All sets of amplicons were benchmarked with the mock community. In addition, we assessed the impact of using different databases for taxonomic assignment as there has been limited work assessing the importance of the reference databases for StrainID (11, 15).

## Materials and Methods

### Biospecimen Collection

Saliva was collected from subjects, birth to ≤21 years of age, enrolled in a prospective study at a single center between March 2021 and August 2022 (IRB approval #21-004). The study’s primary aim was to identify a biosignature of diagnostic value for multisystem inflammatory syndrome in children (MIS-C) (30). Saliva was collected at time of enrollment using the SalivaBio Infant’s Swab Collection kit for subjects birth to <6 years of age (5001.8;5001.5, Salimetrics_®_, PA, USA) or the SalivaBio Passive Drool Collection Aid (5016.04, Salimetrics, PA, USA) for subjects >6 years of age.

Mouse fecal pellets were collected from individually housed mice belonging to one of two strains: Wild-type C57BL/6 strain, and a BTBR autism model mouse line with mitochondria from the C57BL/6 wild type strain (BTBR-mt^B6^) (31). Mice were cared for as described previously (32). Briefly, mice were kept in the same room and received the same food. Human fecal samples were donated by undergraduate students from the University of Connecticut (IRB approval H14-253). Samples were collected using the OMNIgene®•GUT Sample Collection Tube (OM-200, DNAgenotek™, Ontario, Canada). All sample types were stored at -80°C until use.

### Initial Processing of Saliva

Saliva samples were frozen and stored at -80°C for a minimum of 24 hours and then thawed at room temperature for approximately 20 minutes. Debris was pelleted by centrifugation for 15 minutes at 200 rcf and 4°C. A 250 μL aliquot of the supernatant was held at 56°C for 30 minutes to heat-inactivate any present SARS-CoV-2. Samples were treated with 6.25 μL of Proteinase K, which was subsequently inactivated by heating samples to 95°C for 5 minutes. Processed saliva samples were stored at -80°C prior to DNA extraction.

### DNA extraction

DNA was extracted using the Shoreline Complete Lyse & Purify kit (Intus Biosciences, Farmington, CT, United States), which uses an alkaline lysis and magnetic bead purification. Extractions were performed according to the manufacturer’s protocol, with slight differences for each sample type. Saliva samples were thawed on ice, and between 50 μL and 200 µL of saliva was pelleted by centrifugation at 5,500 rcf for 10 min. Pellets were resuspended in 50 μL nuclease-free water, and then extracted. For mouse fecal pellets, approximately ¼ of a pellet was placed in 50 μL nuclease-free water for extraction. For human fecal samples, 10 μL of sample from the OMNIgene®•GUT tube was added to 40 μL nuclease-free water. After extraction, samples were eluted into a total of 40 μL of TE buffer. To confirm successful extraction of bacterial DNA, samples were screened by qPCR targeting the 16S rRNA gene.

### PCR amplification and library prep

Extracted DNA was amplified with the Intus Biosciences Amplify kits according to the manufacturer’s protocol and as previously described (12). Briefly, 10 µL of DNA and 10 µL of PCR mix (Intus Biosciences, Farmington, CT, United States) were added to each well with barcoded primers. The StrainID Amplify kit was used for all sample types. Additionally, the saliva samples were amplified with the V1-V3 Wave kit (Intus Biosciences, Farmington, CT, United States), and all fecal samples were amplified with the V4 Wave kit (Intus Biosciences, Farmington, CT, United States). As a control and to test for primer bias, we also amplified a ZymoBiomics Mock DNA community (Catalog number D6305, Zymo Research, Irvine, CA, United States) with each amplicon type. This mock community contained equal concentrations of genomic DNA from 8 diverse species of bacteria (*Listeria monocytogenes*, *Pseudomonas aeruginosa*, *Bacillus subtilis*, *Escherichia coli*, *Salmonella enterica*, *Lactobacillus fermentum*, *Enterococcus faecalis*, *Staphylococcus aureus*) representing a wide variety of GC content to help identify bias. Amplicons were screened on a QIAxcel Advanced system (Qiagen, Germantown, MD, United States) using the Fast Analysis protocol. Samples were normalized by pooling 5 μL per sample together based on band intensity. Dark bands indicating high levels of amplification formed the first pool, and light bands indicating lower levels of amplification formed the second pool. Both pools were cleaned using the GeneRead Size Selection Kit (Qiagen, Germantown, MD, United States) according to the manufacturer’s protocol and resuspended in 50 µL of elution buffer. After verifying the purity of both initial pools, amplicons were combined and sent for sequencing. StrainID samples were sequenced on a PacBio Sequel IIe, while the V1-V3 and V4 samples were sequenced on an Illumina MiSeq. For the V4 amplicon, we used the v2 2×250 kit, and for theV1-V3 amplicon, we used the v3 2×300 kit.

### Data processing and statistical analysis

Raw sequences were first demultiplexed with Illumina BaseSpace for the V1-V3 and V4 amplicons, and SBAnalyzer for the StrainID amplicons. To account for any biases from the primers, we generated *in silico* reads by trimming the StrainID reads based on the V1-V3, V4, and full 16S primer sequences. Reads were error corrected with the software package DADA2 to generate exact sequences known as amplicon sequence variants (ASVs) (33). Taxonomic assignment was performed with the Athena database (12) and the publicly available Greengenes2 database (2024.09) (34). Data analysis was performed in QIIME2 (2024.5) and R (35). To account for unequal read depth between methods, fecal samples were rarefied to an even depth of 5,716 reads per sample, and saliva samples were rarefied to 7,026 reads. For phylogenetic analyses, the QIIME2 implementations of mafft and fasttree2 were used to generate a tree (36, 37). PcoA plots were visualized with microViz (38), and box plots were created with ggplot2 (39).

Using the core-metrics-phylogenetic function from the diversity plug-in in QIIME2, beta diversity was calculated for all samples by Weighted UniFrac, Unweighted UniFrac, Bray-Curtis, and Jaccard, which differ in how they account for phylogenetic relationships and relative abundances of taxa (35). Bray-Curtis accounts for relative abundance but not phylogenetic relationships; Jaccard ignores phylogeny and is based on presence or absence; Weighted UniFrac accounts for both phylogeny and relative abundance, while Unweighted UniFrac is a phylogenetic presence-absence metric. Differences between groups of samples were assessed using PERMANOVA calculated with the adonis package in QIIME2, which generated R^2^ and adjusted p-values (40). To compare differences in the relative abundances and taxonomic classifications between sequencing methods, descriptive statistics were calculated with ggpubr’s desc_statby tool, and significance was assessed with ggpubr’s compare_means tool (41). The code used is available at https://github.com/joerggraflab/Code-for-OSullivan-2025.

## Results

### Use of a mock DNA community to evaluate StrainID accuracy

To test for any biases introduced by the StrainID primers, we evaluated a mock DNA community of eight taxa against the expected relative abundance. The mock community was amplified and sequenced five times across four sequencing runs. Across all sequencing runs, all eight genera were identified consistently at approximately the correct relative abundances reported by the manufacturer. The largest discrepancies were an overrepresentation of *Bacillus* (mean 0.279 ± 0.057 observed vs 0.174 expected) and an underrepresentation of *Staphylococcus* (mean 0.0727 ± 0.0121 observed vs 0.155 expected) (Figure 2A). Additional analyses using a single replicate of the DNA mock community for the other amplicons also were performed (Supplemental Figure 1). The StrainID ASVs were bioinformatically truncated to correspond to full-length 16S, V1-3 and V4 and received the prefix *in silico*. Both *in silico* amplicons derived from StrainID reads closely resembled the community structure with whole length StrainID, indicating that differences relative to the theoretical values are driven by primer bias. Notably, the V1-V3 amplicon data was in poor agreement with the theoretical composition, with *Lactobacillus* and *Staphylococcus* each representing approximately 1% of the observed community (Supplemental Figure 1).

**Figure 2:**
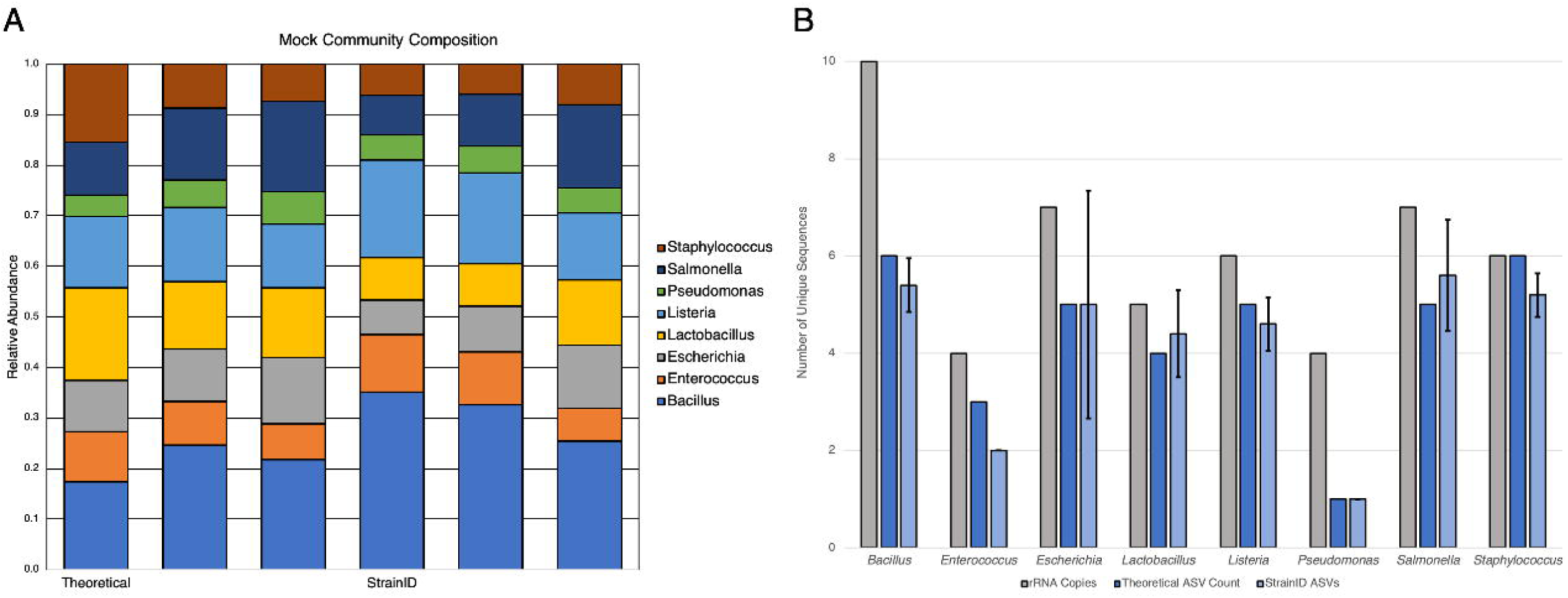
Analysis of a mock community with StrainID. (A) The composition of a defined DNA community against the predicted composition, shown with genus-level classifications. The mock community was amplified five separate times and sequenced across four different runs. (B) Mean number of ASVs for each taxa, compared to the expected number of ASVs and total rRNA operon copies in the reference genomes of all 8 members of the mock community. Error bars represent 1 standard deviation.

Each member of the mock community had multiple copies of the rRNA operon in its genome, with as many as 10 copies in *Bacillus subtilis.* Sequences including more of the operon capture greater sequence diversity between copies. As such, longer amplicons would be expected to result in additional ASVs for each taxon. Using the published genomes for the strains in the mock community, we identified the number of rRNA copies and how many unique sequences were present over the length of the StrainID amplicon, establishing a theoretical ASV count to compare against (Figure 2B). Comparisons to shorter amplicons demonstrated that as sequence length increased, the number of ASVs predicted and observed increased (Supplemental Figure 2). The number of ASVs observed with StrainID closely matched the predicted number, which is higher than the ASVs for V1-V3 and V4 (Figure 2B, Supplemental Figure 2).

### Phylum-level Classification of Bacterial Communities

After confirming the accuracy of StrainID with the mock community, we sought to compare its performance to SRA with salivary and fecal samples at higher taxonomic levels. Included in the comparison were 46 saliva samples, eight human fecal samples, and fecal samples from six wild type C57BL/6 strain mice and nine samples from a strain of BTBR strain complemented with mitochondria from the wild type strain (BTBR-mt^B6^) mice (31).

The overall compositions of the microbial communities were compared between the different amplicons. At the phylum level with classification using the Athena database, differences were observed between V1-V3 and StrainID in the saliva samples. This was not unexpected based on the observed primer bias in the mock community. Although Firmicutes were the most dominant phylum with both methods, their overall abundance was higher with StrainID compared to V1-V3 (median abundance of 72.8% vs 62.0%, respectively, p<0.05) (Figure 3A). The relative abundances of Actinobacteria, Proteobacteria, and Saccharibacteria also differed significantly between V1-V3 and StrainID (p<0.05). Notably, Saccharibacteria was not detected with StrainID. Analysis of the genome for an oral isolate of Saccharibacteria, *Nanosynbacter lyticus* TM7x (CP040011.1), revealed that the StrainID reverse primer had several mismatches, and may fail to bind (42). Further, in these taxa the operon is non-contiguous due to the presence of *pyrD* and a gene encoding a hypothetical protein between the 16S and 23S rRNA genes. The resulting 7,069 bp amplicon is likely too long for the amplification conditions used and/or would be discarded during standard bioinformatic quality control steps. Fecal samples from both humans and mice were dominated by Firmicutes and Bacteroidetes, followed by lower abundances of Actinobacteria and Proteobacteria with both V4 and StrainID (Figure 3B). In total, these results suggest minimal differences in amplification in the fecal samples, but potentially biologically relevant differences within the saliva samples.

**Figure 3:**
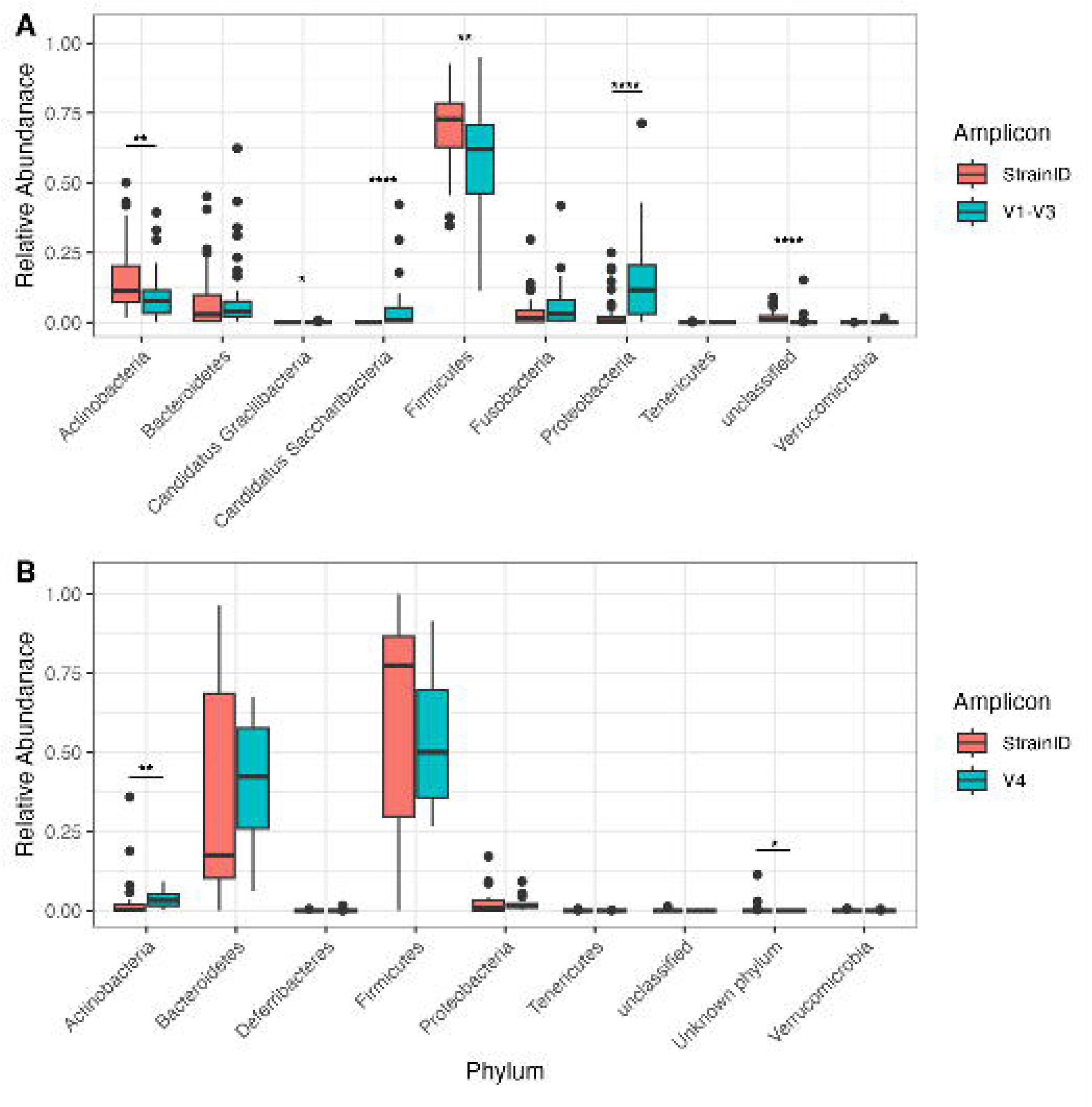
Phylum-level abundances of saliva and fecal samples. Box and whisker plots of the relative abundances of the phyla observed in saliva (A) and fecal (B) samples. Horizontal bars represent median abundances. (*p<0.05, **p<0.005, ****p<0.00005)

### Taxonomic Classification Improves with StrainID

Performance of read classification at different taxonomic levels was evaluated with the reference databases Athena and Greengenes2. The initial analysis was performed with the Athena database. For saliva samples, *in silico* 16S performed the best at the genus level, with a median 100% of reads classified per sample. V1-V3 and StrainID also performed well (median 99.7% and 98.8%, respectively), though differences in taxonomic assignment were significantly different between each group (p≤0.005) (Figure 4A). At the species level, *in silico* 16S still resulted in the most reads classified (median 99.2%), though V1-V3 (median 92.5%) and StrainID (median 89.4%) no longer differed significantly. None of the amplicons differed significantly from each other for classification at the strain level (median classification between 20.8% and 22.2%).

**Figure 4:**
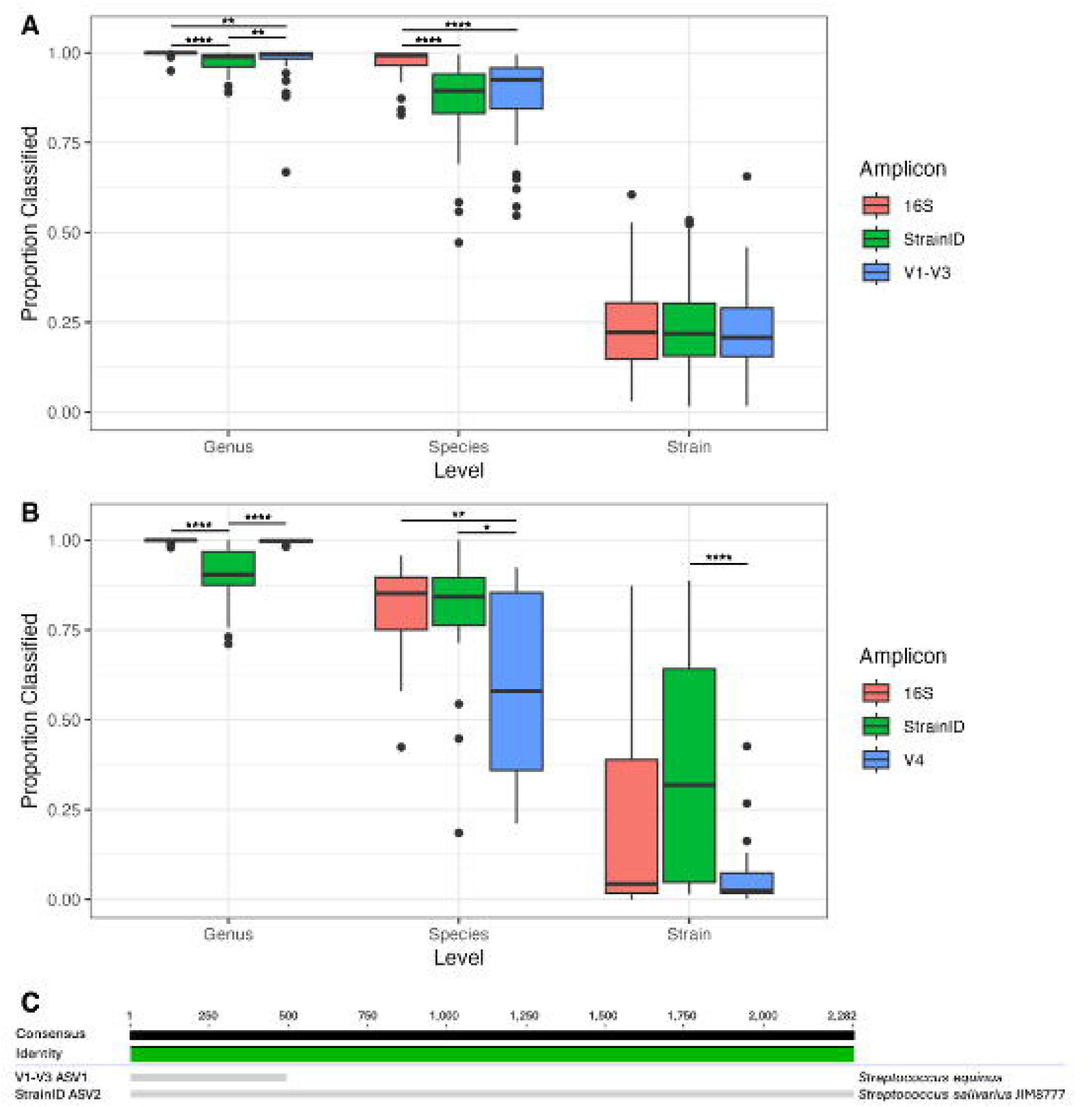
Read classification by amplicon. Box and whisker plots for the proportion of reads classified at the genus, species, and strain levels with the Athena database for both saliva (A) and fecal samples (B). Horizontal bars represent the median. (C) Alignment of the most abundant ASV overall in saliva samples with both StrainID and V1-V3, with the highest level of taxonomic assignment provided with the Athena database. (*p<0.05, **p<0.005, ****p<0.00005)

Because Greengenes2 is a 16S rRNA database and does not include ITS and 23S rRNA sequences, StrainID reads are inherently incompatible, unless first trimmed to include only the 16S rRNA region for classification. Furthermore, the shorter length means the database does not support strain-level classifications. Here, genus level classification performed similarly across all methodologies, though species-level classifications were more successful with SRA (Supplemental Figure 3). This is explained in part by the Greengenes2 containing a mix of full length and partial 16S rRNA sequences, meaning there are more potential matches for shorter ASVs.

For fecal samples, a different pattern was observed than that found in saliva using the Athena database. At the genus level, *in silico* 16S and V4 resulted in significantly higher levels of read classification than StrainID (median 90.5%, p<0.0005) but did not differ significantly from each other (median 100% and 99.8%, respectively) (Figure 4B). With species-level classifications, however, the benefit of longer reads became clear. V4 performed significantly worse than the other amplicons, classifying a median 57.9% of reads compared to 85.2% and 84.3% for *in silico* 16S and StrainID, respectively (p<0.05). Finally, for strain-level classifications, StrainID significantly outperformed V4 (median 31.9% vs. 2.5%, p<0.0005). Although other comparisons were not significant, StrainID did result in more strain-level classifications than did *in silico* 16S did (median 4.3%).

The most abundant ASV present in human saliva samples with StrainID was identified with Athena as *Streptococcus salivarius*, a commensal microbe commonly found in the human oral cavity. The most abundant ASV with V1-V3 was identical to the StrainID ASV for the overlapping regions. Despite this, it was identified instead as *S. equinus*, which as the name implies, is associated primarily with horses (Figure 4C). Given the sample type, the classification of *S. salivarius* seems far more likely to be correct. With Greengenes2, this ASV was identified only as *Streptococcus*, even with the whole length 16S sequence, demonstrating the benefit to taxonomic resolution provided by the inclusion of the ITS and 23S rRNA regions.

### StrainID increases statistical power of beta diversity metrics

To assess the practical application of the different approaches, we next sought to test the ability to differentiate between groups of samples by four different beta-diversity metrics: Bray-Curtis, which accounts for relative abundance but not phylogenetic relationships; Jaccard, which ignores phylogeny and is based on presence and absence; Weighted UniFrac, which accounts for both phylogeny and relative abundance; and Unweighted UniFrac, which is phylogenetic presence-absence metric. The saliva samples were collected using two methods, the passive drool and swab methods. We assessed if there were differences in the salivary microbiome depending on the collection method using PERMANOVA calculated with the adonis tool in QIIME2. The largest amount of variation explained by the collection method was with the StrainID amplicon and weighted UniFrac (R^2^=0.064, p=0.002) (Figure 5A). With the V1-V3 primer set, this comparison was both not significant and explained less variation (R^2^=0.021, p=0.471) (Figure 5B). The *in silico* approaches performed similarly to StrainID though the differences between collection methods were slightly less significant (16S R^2^=0.058, P=0.018; V1-V3 R^2^=0.062, P=0.011) (Figure 5C-D). All other comparisons were significant (p<0.05) and explained between 2.5% and 5.5% of variation between samples (Table 1).

**Figure 5:**
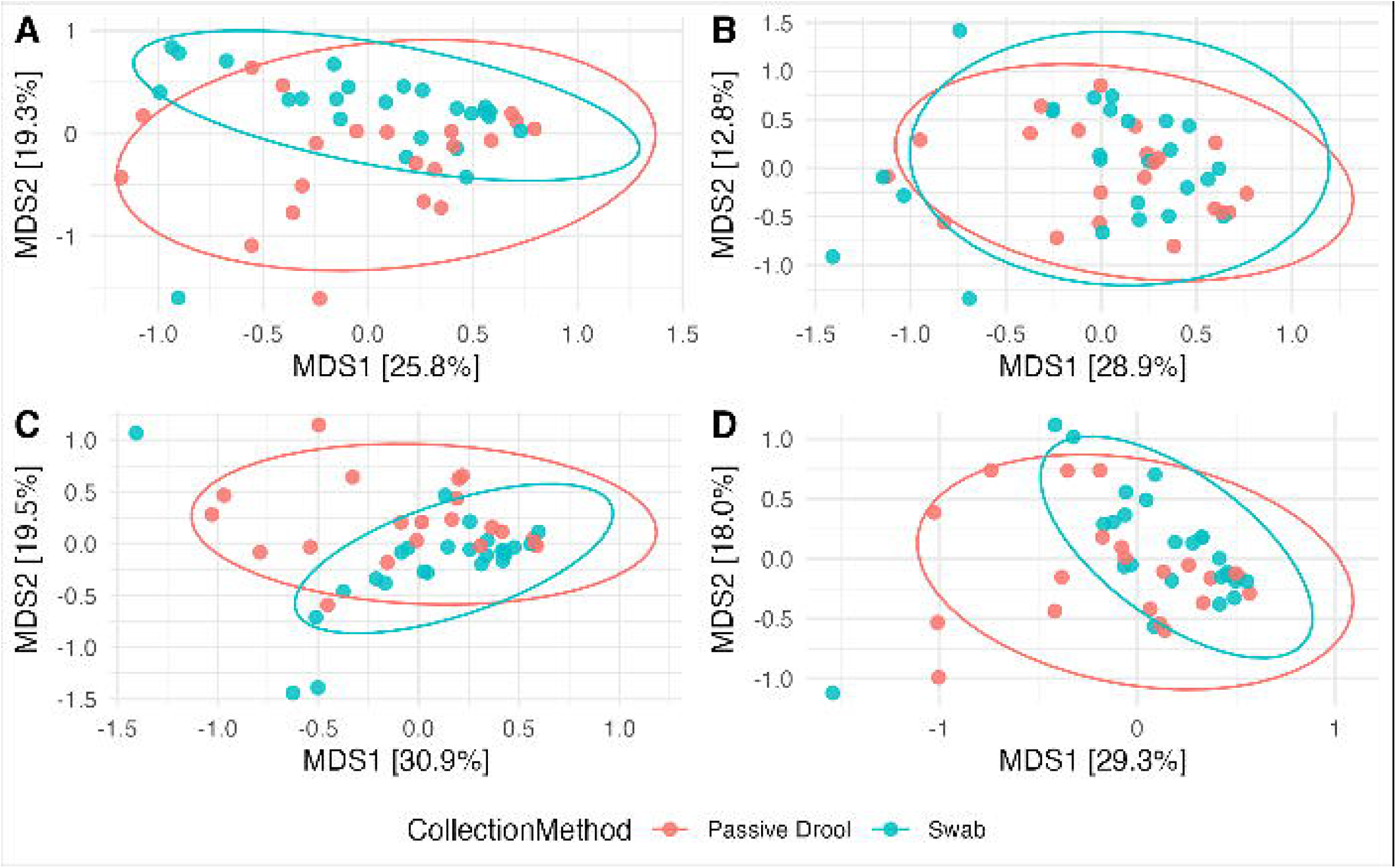
PCoA of Weighted UniFrac for saliva samples. PCoA plots of StrainID (A), V1-V3 (B), *in silico* 16S (C), and *in silico* V1-V3 (D). Samples were grouped by the saliva collection method, and confidence intervals are denoted by ellipses.

**Table 1:**
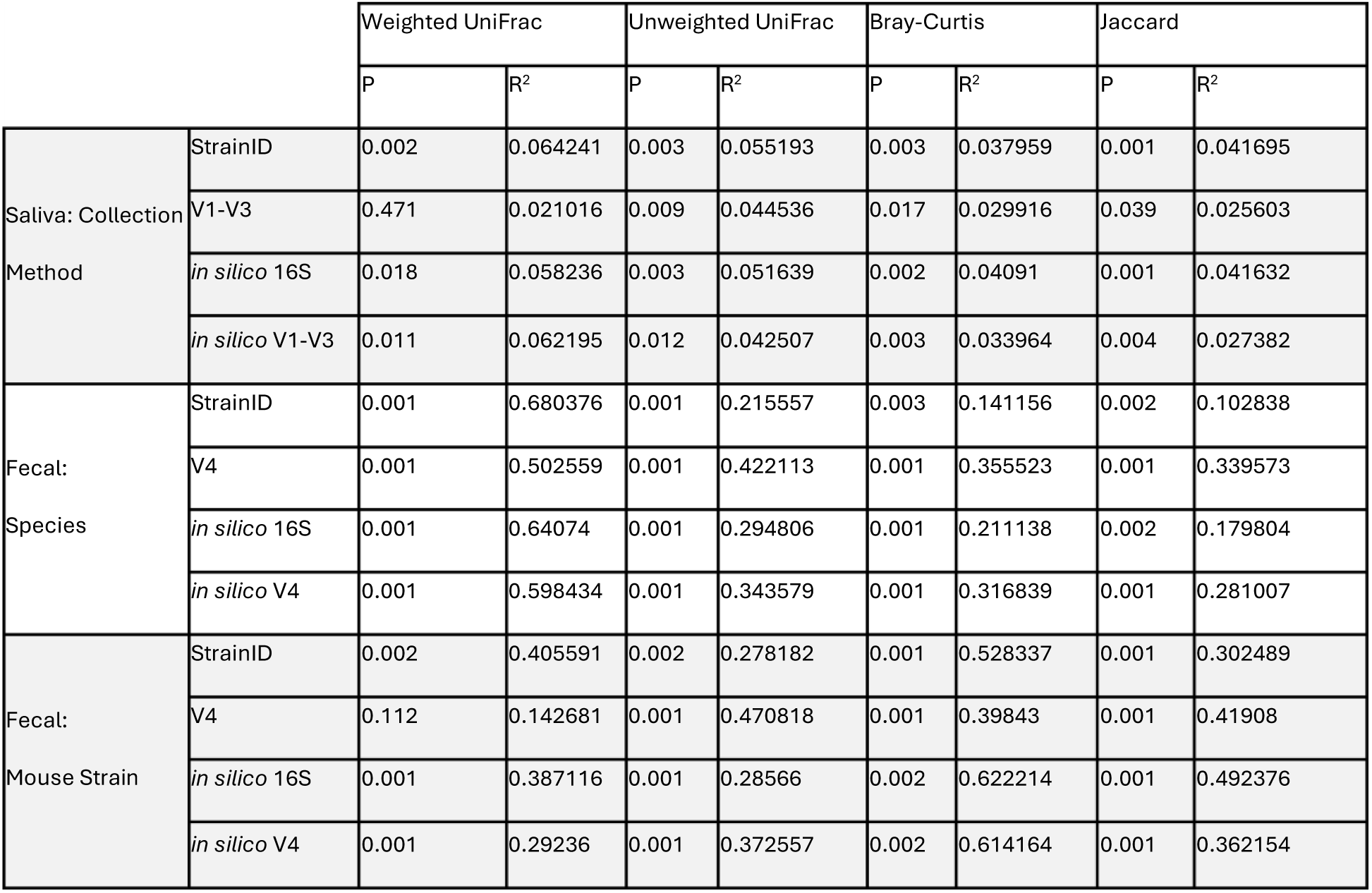
Adonis PERMANOVA of Saliva and Fecal Samples.

For fecal samples, two comparisons were performed: First, by host species (human vs. mouse), followed by strain of mouse (C57B/L6 vs BTBR-mt^B6^). Samples differed significantly by species by all four beta-diversity metrics and for all primer sets (Table 1). The most variation was explained by host species with weighted UniFrac, where StrainID outperformed V4 (R^2^=0.680, p=0.001 and R^2^=0.502, p=0.001, respectively) (Figure 6A-D, Table 1). When comparing mouse strains, samples differed significantly and explained between 27.8% and 61.4% of variation for all primer pairs and all metrics except with weighted UniFrac. With the weighted UniFrac metric, the strains did not differ significantly with the V4 primers (R^2^=0.143, p=0.112) (Figure 6F). For the same comparison, StrainID was highly explanatory (R^2^=0.406) and highly significant (p=0.002) (Figure 6E). The *in silico* approaches based on the StrainID reads were also highly significant (p=0.001) but explained less variation (16S R^2^=0.387, V4 R^2^=0.292), suggesting that at least some of the difference was due to primers used for amplification (Figure 6G-H).

**Figure 6:**
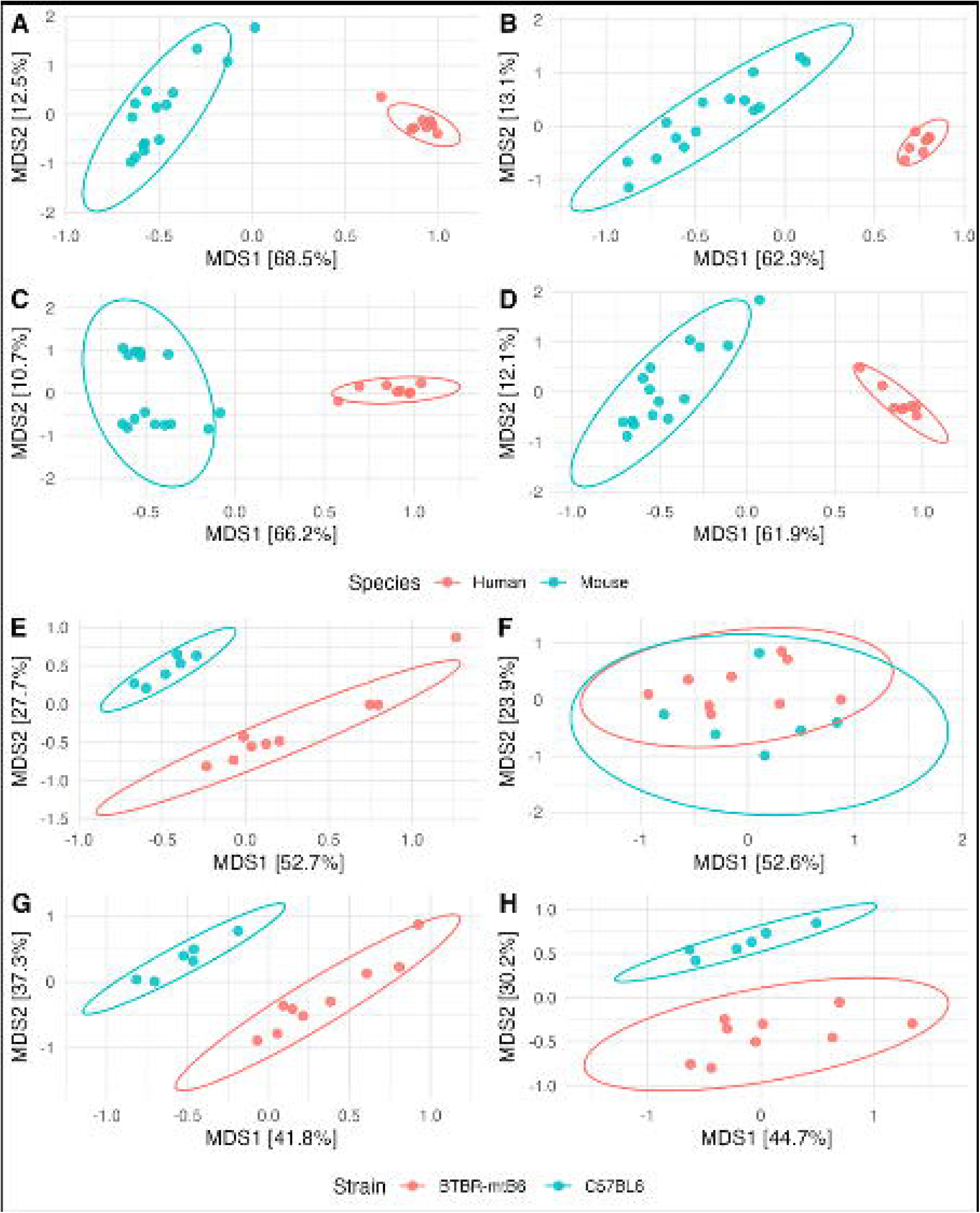
PCoA of Weighted UniFrac for fecal samples. (A-D) PCoA plots of all fecal samples with StrainID (A), V4 (B), *in silico* 16S (C), and *in silico* V4 (D). Samples were grouped by the host species, and confidence intervals are denoted by ellipses. (E-H) PCoA plots of mouse samples with StrainID (E), V4 (F), *in silico* 16S (G), and *in silico* V4 (H). Samples were grouped by the mouse strain, and confidence intervals are denoted by ellipses.

## Discussion

Overall, these results validate StrainID for use with human saliva samples in microbiome sequencing studies, not only by improving the ability to differentiate distinct sample types or cohorts, but also by providing enhanced taxonomic insights. Additionally, they further establish StrainID as a robust methodology that compares favorably to traditional SRA approaches. In terms of taxonomic accuracy, StrainID exceeds the performance of V1-V3 for saliva samples, and is similar in performance to V4 for fecal samples, albeit with some primer bias introduced.

However, all primer sets introduce some bias (43, 44) and the ability to use one set of primers with excellent taxonomic resolution for all communities is another advantage of StrainID. The amount of primer bias seen with V1-V3 exceeded that observed with StrainID, suggesting that StrainID may be more accurate. There is no loss in performance when comparing communities of samples with StrainID compared to SRA methods. In some cases, StrainID was actually more effective than traditional approaches at distinguishing groups, especially with phylogenetic metrics.

StrainID also provides the benefit of longer read length which increased the accuracy of taxonomic assignment. Database selection is an important part of taxonomic classification and can significantly impact results (43, 44). Databases containing longer sequences are better able to provide matches with a high resolution but contain fewer sequences, which means certain taxa may be poorly represented, especially from understudied environments. With the Athena database, short read methods could slightly outperform StrainID. One potential explanation is that the number of sequences from bacteria found in the oral cavity in the Athena database is relatively small. This could result in fewer good matches over the whole length of StrainID. Meanwhile, the database is large enough for SRA reads to match, but small enough that there will not be multiple matches to different taxa which would result in those hits being discarded. We also found that for a small proportion of ASVs, StrainID can fail to identify taxa even at the genus level more often than observed with *in silico* 16S approaches. These differences are likely driven by mismatches over the ITS region due to the failure of existing databases to capture the true diversity of sequences present in nature. For ASVs not identified at the genus level, trimming the sequences to just the whole length 16S rRNA region and reclassifying should, at a minimum, enable a coarser classification.

In addition, one can use the sequence information for the design of diagnostic primers or probes for fluorescent *in situ* hybridization. As the number of genomes in the public databases continuously increase, the accuracy of reference databases designed for long-reads should increase as well and allow for even more sequences to be identified at the species level. When specifically comparing taxonomic assignment of saliva samples between V1-V3 and StrainID, we found that V1-V3 occasionally outperformed StrainID. However, while these differences were statistically significant, they were small enough that they likely are not biologically meaningful.

StrainID is not without shortcomings, which must be considered when selecting primers. For one, the amplicon design will fail to identify any bacteria with non-continuous rRNA operons, such as in *Helicobacter*, in which the 16S rRNA gene is not linked to the 23S and 5S rRNA genes (45). This was seen in our dataset with the phylum Saccharibacteria found in saliva only with SRA due to incompatibility with StrainID. Saccharibacteria is a potentially significant phylum for saliva, and past studies have correlated this taxon with disease states (46). However, all known Saccharibacteria are episymbionts of other bacteria (42). As such, it is possible that any signal associated with specific species of Saccharibacteria could be identified by their symbionts, which likely would be compatible with StrainID. That StrainID will result in the loss of some taxa is a significant limitation of the approach and must be balanced against the benefits of improved resolution.

Additionally, the long sequence length of StrainID can lead to a low percentage of nonunique sequences. Because DADA2 relies on repeat observations, this can lead to excess reads being lost unless “pooling” the sequences from multiple samples together is done, which greatly increases the required computational time and power. As a result, Intus Biosciences has begun moving away from DADA2 and has started to develop a proprietary alternative software package for use with StrainID. Additionally, at the time of this publication, the cost per sample with StrainID is typically higher than with SRA methods due to limits in multiplexing. SRA barcodes support up to 384-plex sequencing runs. To date, StrainID is limited to a maximum of 112 unique barcodes. Further development and verification of additional barcodes for StrainID could help minimize this weakness. Adapting the primers to be compatible with PacBio’s Kinnex approach, which concatenates multiple amplicons together in a single molecule, could further assist in increasing the number of samples per run, in addition to increasing the average read depth, bringing the costs of StrainID more in-line with SRA sequencing.

StrainID presents several important strengths. Our results suggest that StrainID improves taxonomic resolution and assignment accuracy when studying the microbiome. This is especially important for detecting differences in microbiome composition between different cohorts in clinical studies. Additionally, rRNA fingerprinting with StrainID allows identification of ribotypes (12). This is especially powerful in studies where tracking an individual ribotype would be useful, such as tracking transmission, outbreak, or temporal studies where continued presence of a ribotype would suggest its stability as a community member (47).

In conclusion, StrainID is accurate, matches or exceeds the performance of SRA, and improves taxonomic resolution. Given the advantages of this methodology, long read sequencing approaches should be given consideration when designing a study where resolution is especially important. In recent years, there has been a greater shift towards the use of metagenomics in microbiome research due to the completeness of the data it provides compared to SRA. However, whole 16S rRNA sequencing is also becoming more common, especially as Oxford Nanopore’s accuracy continues to increase and PacBio’s cost per sample decreases. StrainID can be used in lieu of whole length 16S rRNA sequencing to increase resolution for a similar cost and represents a more powerful approach for microbiome studies.

## Acknowledgements

Research reported in this publication was supported by the Eunice Kennedy Shriver National Institute of Child Health and Human Development of the National Institutes of Health under award numbers R61HD105613 and R33HD105613. We would like to express our immense gratitude to Robert and Francine Shanfield for their generous donation in support of this research, without which this article would not be possible. We would also like to thank our colleagues Drew Bidmead, Adriana Camacho, Carlie DeFelice, Kristina Dibble, Kemar Edwards, Hilda Giraldo, Rebecca Hibbard, Kelly Howard, Carson Karanian, Karen Kulas, Callista Lajeune, Stephanie Lesmes, Makayla Murphy, Isabel Orbe, Celina Porcaro, Rebecca Radolf, Rosa Rodrigues, Jessika Rodrígueza, Noah Schulman, and Beatriz Vanegas for their vital contribution to the project, including participant recruitment, biospecimen collection, processing and testing, and data capture. Mouse Fecal samples were obtained from the Lawrence Lab with IACUC approval (protocol # 21-278) received on 07/19/2021. The Wadsworth Center maintains a PHS-assured and AAALAC-accredited animal use program (Animal Welfare Assurance Number D16-00115). Sequencing was performed by the University of Connecticut Microbial Analysis and Resource Services, the University of Delaware Sequencing and Genotyping Center, and the University of Florida Interdisciplinary Center for Biotechnology Research.

We also thank Mark Driscoll and Eric Jackson from Intus Biosciences for their advice and assistance with the StrainID data analysis.

## Data availability

Raw read data are available in the NCBI SRA database under project ID PRJNA1314305. Additional reads used can be found under project ID PRJNA910511 under biosample IDs SAMN32127837 through SAMN32127842 and SAMN32127844 through SAMN32127851.

---

**Supplemental Figure 1:**
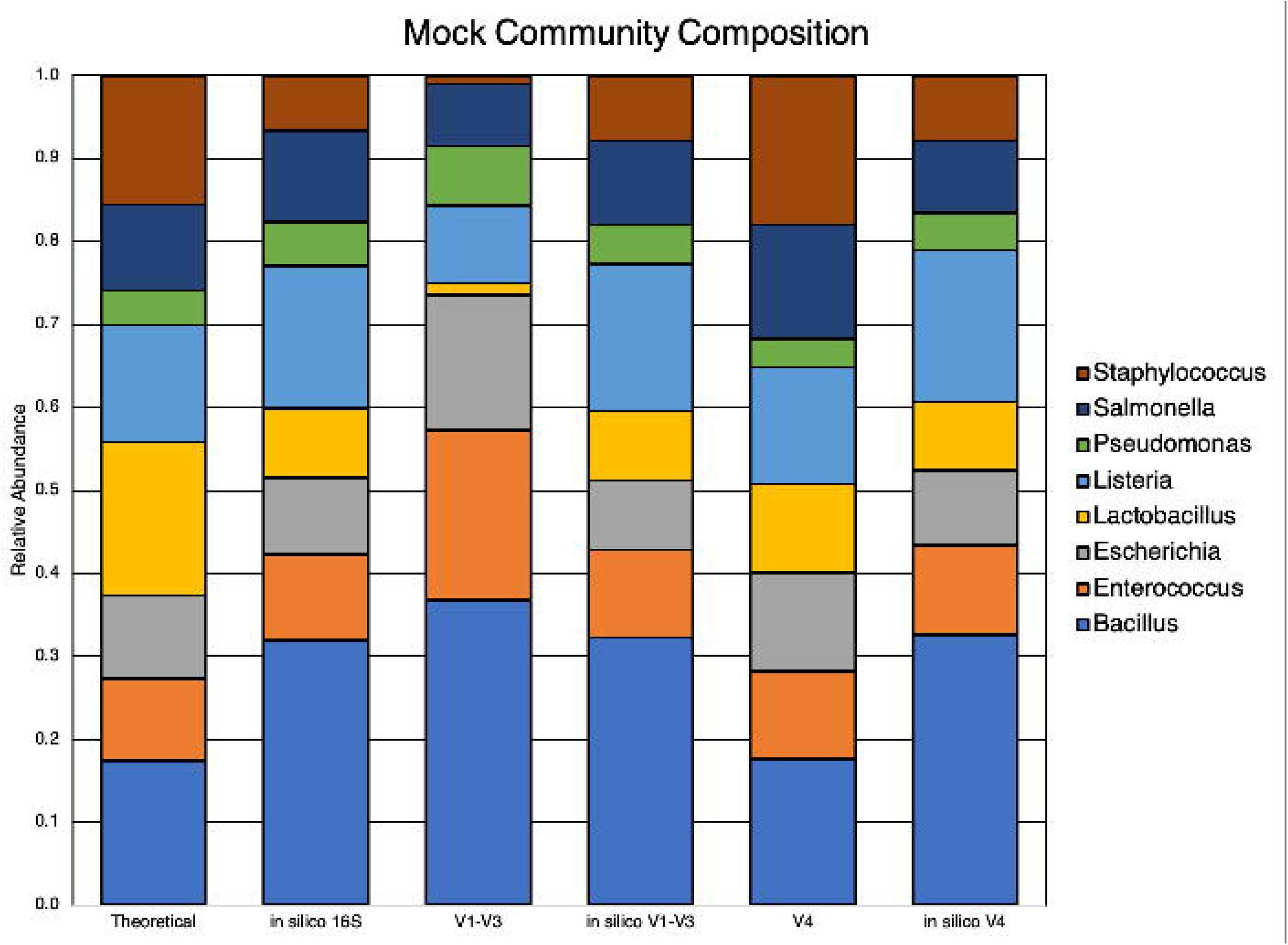
The composition of a defined DNA community against the predicted composition, shown with genus-level classifications for each amplicon type tested, excluding StrainID. The *in silico* amplicons were derived from StrainID reads that were trimmed to the appropriate length prior to ASV calling.

**Supplemental Figure 2:**
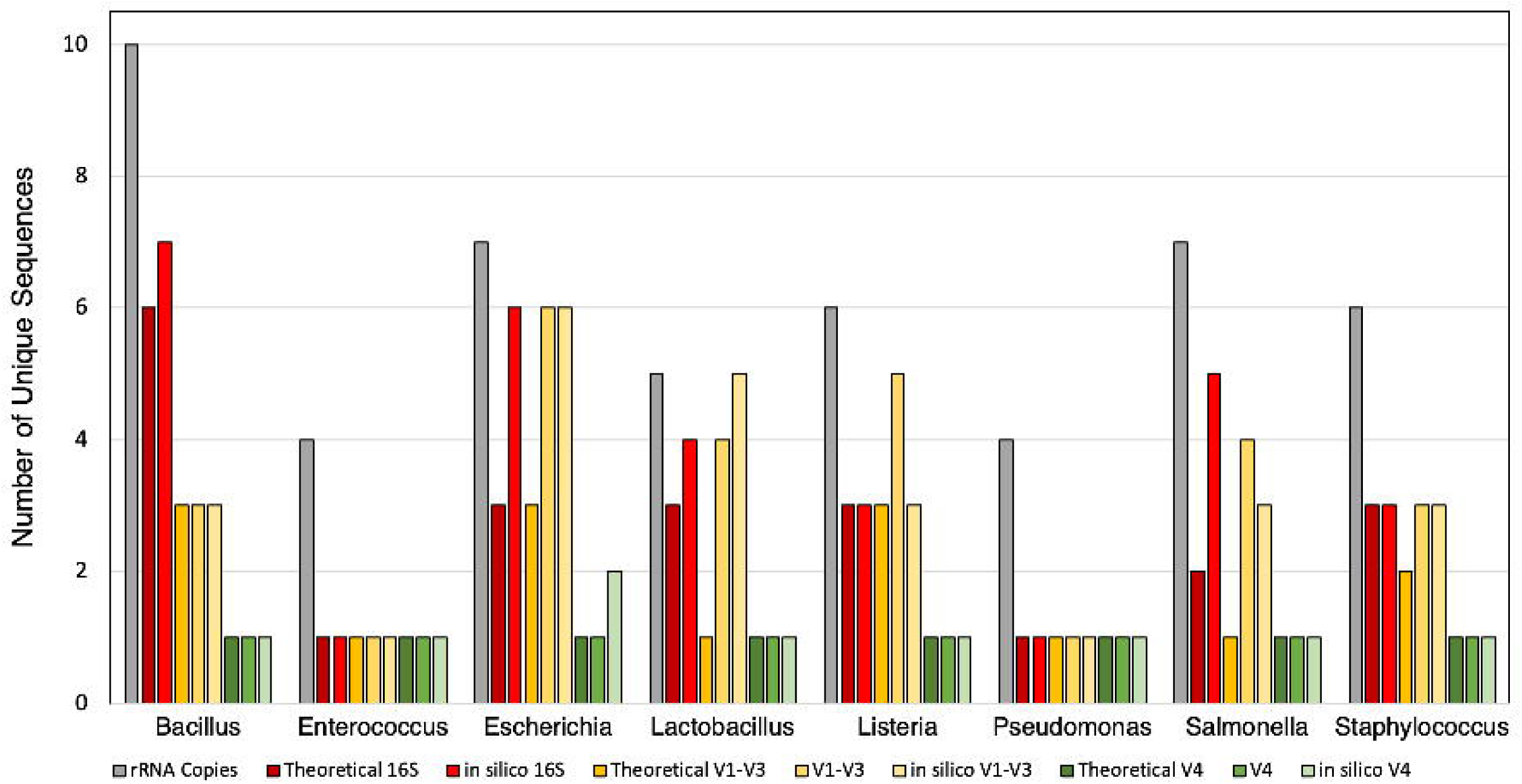
The number of ASVs for each taxa, compared to the expected number of ASVs and total rRNA operon copies in the reference genomes of all 8 members of the mock community for all amplicon types tested.

**Supplemental Figure 3:**
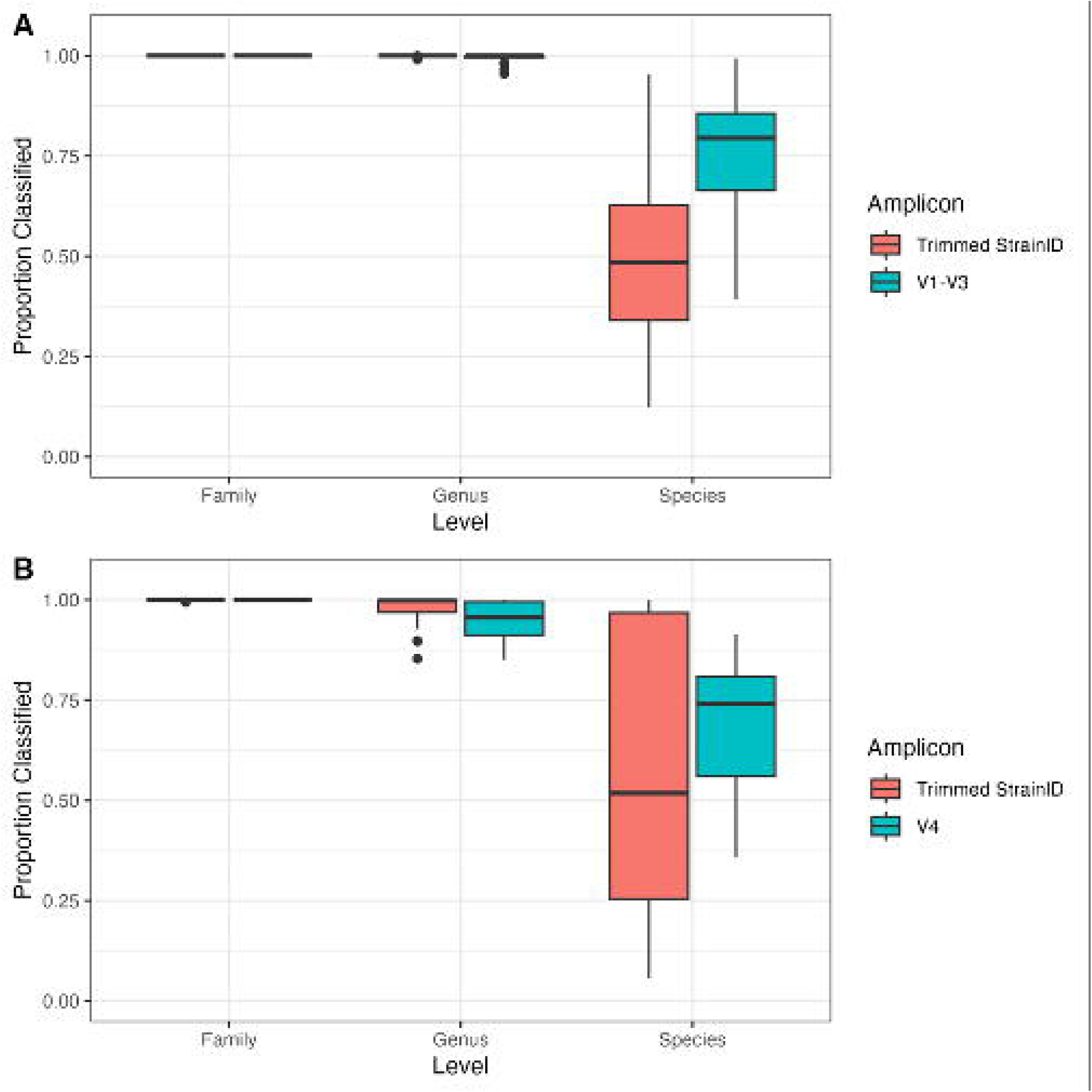
Box and whisker plots for the proportion of reads classified at the family, genus, and species levels with the Greengenes2 database for both saliva (A) and fecal samples (B). StrainID ASVs were trimmed to include only the full-length 16S region to be compatible with the database.

